# Nuclear organisation and replication timing are coupled through RIF1-PP1 interaction

**DOI:** 10.1101/812156

**Authors:** Stefano Gnan, Ilya M. Flyamer, Kyle N. Klein, Eleonora Castelli, Alexander Rapp, Andreas Maiser, Naiming Chen, Patrick Weber, Elin Enervald, M. Cristina Cardoso, Wendy A. Bickmore, David M. Gilbert, Sara C. B. Buonomo

**Author notes:** These authors contributed equally to the work (Ilya M. Flyamer, Kyle N. Klein).

## Abstract

Three-dimensional genome organisation and replication timing are known to be correlated, however, it remains unknown whether nuclear architecture overall plays an instructive role in the replication-timing program and, if so, how. Here we demonstrate that RIF1 is a molecular hub that co-regulates both processes. Both nuclear organisation and replication timing depend upon the interaction between RIF1 and PP1. However, whereas nuclear architecture requires the full complement of RIF1 and its interaction with PP1, replication timing is not sensitive to RIF1 dosage. RIF1’s role in replication timing also extends beyond its interaction with PP1. Availing of this separation-of-function approach, we have therefore identified in RIF1 dual function the molecular bases of the co-dependency of the replication-timing program and nuclear architecture.

## Introduction

In eukaryotes, origins of DNA replication are not activated all at once. Origin firing follows a cell-type specific temporal program known as DNA replication timing. The replication-timing program is mirrored by the spatial distribution in the nucleus of replication foci, which are clusters of about 5 simultaneously activated bidirectional replication forks ^1^. Both spatial and temporal replication patterns are re-established every cell cycle in G1, at the timing decision point (TDP) ^2^, that coincides with chromosomal territories achieving their stable, reciprocal position ^3^ and the re-establishment of chromatin architecture and interphase-nuclear configuration ^4,5^. The spatial organisation of DNA replication is evident at multiple levels. The units of DNA replication timing, replication domains (RD), coincide with one of the basic units of three-dimensional (3D) genome organisation, the topologically associated domains (TADs) ^6^. Recently, *in cis* elements (early replicating control elements-ERCEs) that can simultaneously influence chromatin looping and replication timing have also been identified ^7^. Moreover, the “assignment” of RDs as early or late replicating (the establishment of the replication-timing program), takes place on a chromosome-domain level, prior to the specification of the active origins of replication ^2^. On a global scale, the early and late replicating genomes overlap with the A and B compartments identified by Chromosome Conformation Capture methods (HiC) ^8–10^ and are segregated in the nuclear interior or the periphery of the nucleus and nucleolus respectively. Finally, a recent study from budding yeast has shown that activation of early origins drives their internalisation ^11^. However, no causal link between the temporal and spatial aspects of DNA replication organisation has been established.

RIF1 is a key genome-wide regulator of replication timing ^12–18^. It is also involved in re-establishing spatial chromatin organisation in the nucleus at G1 ^13^, and in the control of replication foci spatial dynamics ^12^. RIF1 could therefore be a molecular connection between the temporal and spatial organisation of DNA replication in mammalian cells.

The molecular function of RIF1 is still unclear, although it is involved in a variety of functions such as DNA repair ^19–30^, telomere length regulation in yeast ^31–35^, cytokinesis ^36^, epigenetic ^37–41^ and DNA replication-timing control. Mammalian RIF1 (266 kDa) interacts with components of the nuclear lamina ^13,42^, behaving as an integral part of this insoluble nuclear scaffold and chromatin organiser. RIF1 associates with the late replicating genome, forming megabase-long domains called RIF1-associated-domains (RADs) ^13^. It is unknown what directs the association to chromatin, but both the N and C terminus of RIF1 have been shown to be capable of mediating interaction with DNA ^33,43–47^. RIF1 has a highly-conserved interaction with Protein Phosphatase 1 (PP1) that is reported to be critical to regulate the firing of individual, late origins of replication ^15,48–51^. Activation of these origins is promoted by RIF1 removal in late S-phase, led by the increasing levels of cyclin-dependent Kinase (CDK) activity ^15,16,48–51^. These studies therefore place the role of the RIF1-PP1 interaction at the stage of execution of the replication-timing program, in S-phase. However, we have also identified a role for RIF1 as a chromatin organiser earlier during the cell cycle, in G1, around the time of the establishment of the replication-timing program ^13^. Rif1 deficiency impacts nuclear architecture, relaxing the constraints that normally limit chromatin interactions between domains with the same replication timing ^13^. It is unknown if Rif1-dependent chromatin architecture establishment affects the replication-timing program, how RIF1 contributes to nuclear organisation, and if and how the interaction with PP1 plays a role in this function. More generally, the functional relationship between nuclear architecture and replication timing is still unclear.

Here, we tackle this question by interfering with the RIF1-PP1 interaction, introducing point mutations in RIF1 that specifically abolish the interaction. Our results show that both replication timing and nuclear organisation depend upon RIF1-PP1 interaction. However, unlike the replication-timing program, we find that nuclear organisation is exquisitely sensitive to RIF1’s dosage. Using this separation-of-function approach, we identify in RIF1 the molecular hub for their co-regulation. In addition, we show for the first time that the replication-timing program can be established and executed independent of the three-dimensional (3D) organisation or the spatial distribution of replication foci.

## Results

### Mouse embryonic stem cells expressing *Rif1*^Δ*PP1*^

RIF1-PP1 interaction promotes the continuous dephosphorylation of MCM4 at replication origins that are “marked” to be activated only during the later part of S-phase ^15,48–51^. This suggests that, through RIF1, PP1 contributes to the control of the timing of firing of individual origins of replication. However, the functional significance of RIF1-PP1 interaction for the establishment and domain-level regulation of the replication-timing program, and in the context of nuclear 3D organisation is unknown.

Mutations that perturb RIF1-PP1 interaction are potential tools to achieve separation-of-function between nuclear organisation and replication timing. We have recently identified the sites within RIF1 that mediate the physical contacts with PP1 (SILK and RVSF motifs, residues 2128–2131 and 2150–2153). Point mutations of these residues reduce PP1 interaction to undetectable levels (RIF1^ΔPP1^: SILK into SAAA and RVSF into RVSA, ^52^). We therefore sought to express the *Rif1*^Δ*PP1*^ mutant in mESCs. However, *Rif1* overexpression is toxic and therefore, in order to create a system to expresses *Rif1*^Δ*PP1*^ in a physiological context, we have utilised *Rif1*^*FH/flox*^ mESCs. In these cells, one allele of *Rif1* contains loxP sites flanking exons 5 to 7 (^19^, *Rif1*^*flox*^)), while the second is a knock-in of a FLAG-HA2 tag (FH) into the *Rif1* locus (*Rif1^FH^*) (^12^, Fig. S1A and B)). We then targeted the FH allele with a mini-gene encoding *Rif1*^Δ*PP1*^. As a control, following the same strategy, we have also knocked-in a *Rif1* wild type mini-gene (*Rif1^TgWT^)*. Thus, Cre-mediated deletion of the *Rif1*^*flox*^ allele leaves either the FH-tagged *Rif1*^Δ*PP1*^, *Rif1*^*TgWT*^ or the parental *Rif1*^*FH*^ allele as the sole source of RIF1, effectively creating inducible FH-tagged *Rif1* hemizygous cells. Upon tamoxifen-mediated Cre recombination, we have then studied the consequences of abolishing RIF1-PP1 interaction in *Rif1*^Δ*PP1/flox*^ (*Rif1*^Δ*PP1/−*^, abbreviated Rif1-ΔPP1), control *Rif1*^*TgWT/flox*^ (*Rif1*^*TgWT/−*^ abbreviated Rif1-TgWT) and the parental *Rif1*^*FH/flox*^ (*Rif1*^*FH/−*^, abbreviated Rif1-FH) cell lines. In agreement with the fact that, upon Cre induction, all the Rif1 FH-tagged alleles are hemizygous, RIF1-ΔPP1, RIF1-TgWT and RIF1-FH, are expressed at comparable levels (Fig. 1A, B and S1C) and RIF1-PP1 interaction is undetectable in RIF1-ΔPP1 (Fig. 1C). Both RIF1-ΔPP1 and RIF1-TgWT have a comparable degree of chromatin-association (Fig. 1D and S1D).

**Figure 1.**
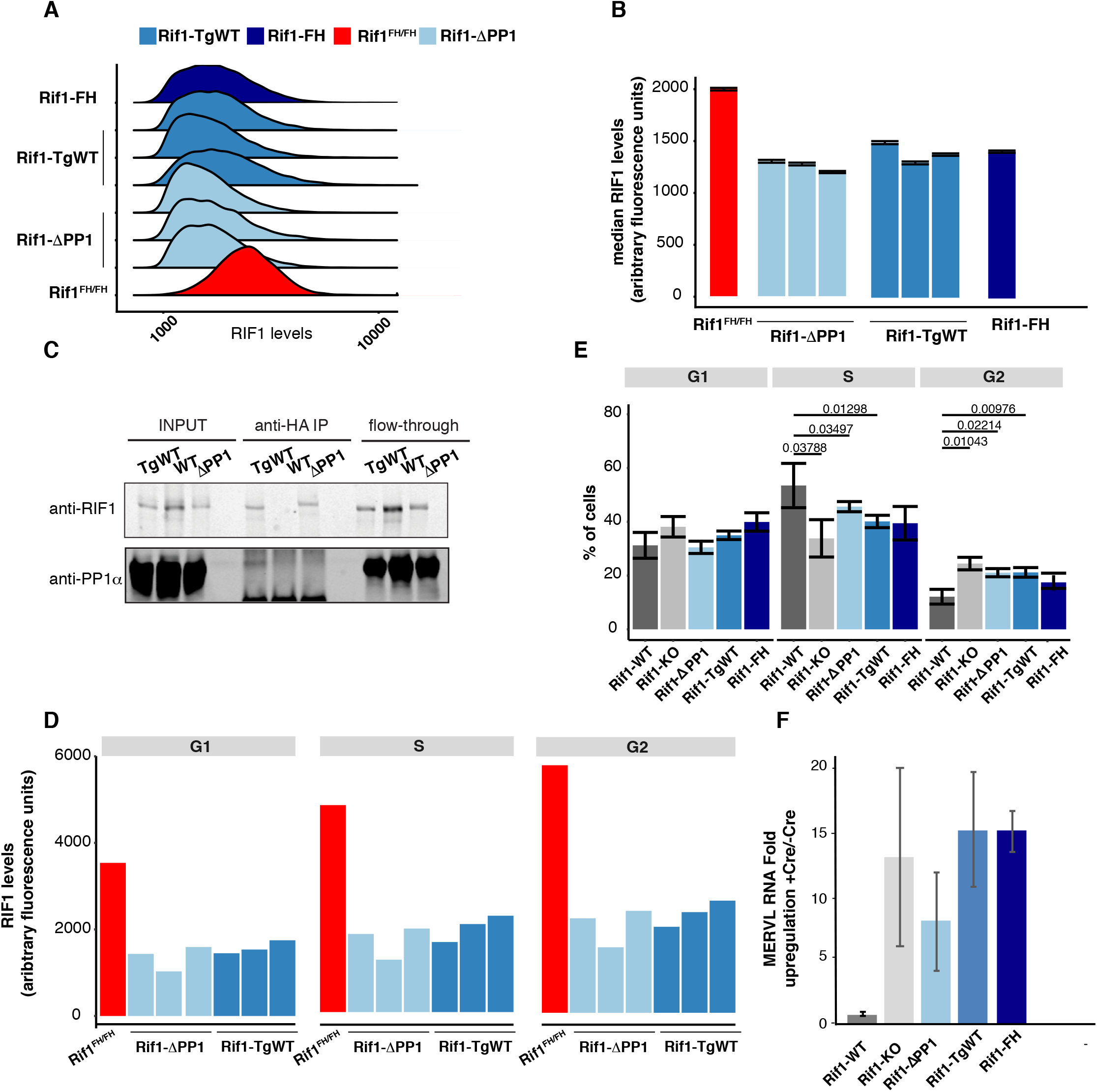
Expression levels and chromatin association of RIF1-TgWT and RIF1-^ΔPP1^ are comparable to those of *Rif1* hemizygous cells. A. Quantitative analysis of total levels of FH-tagged RIF1, measured by intra-cellular FACS staining. Anti-HA mouse ascites 16B12 was used to stain the indicated cell lines. *Rif1*^*FH/FH*^: homozygous knock-in FH-tagged RIF1, as a control of quantitative staining. The plot shows distributions of densities from HA signal, measured in arbitrary units. One representative experiment is shown. B. Quantification from Fig. 1A. The bar plot represents the median intensities for the experiment shown and the error bars indicate 95% confidence intervals. C. Total proteins were extracted from cells expressing either untagged, wild type RIF1 (WT), hemizygote RIF1-TgWT or RIF1-ΔPP1 (HA tagged), and immunoprecipitated with anti-HA antibody. The input, immunoprecipitated complex and flow through were analysed by western blot with anti-mouse RIF1 affinity-purified rabbit polyclonal antibody (1240) and anti-PP1α. D. Quantitative analysis of the levels of chromatin-associated FH-tagged RIF1 throughout the cell cycle (one representative experiment), measured by FACS staining. Cytoplasmic and nucleoplasmic proteins were pre-extracted before fixing chromatin-associated proteins. Anti-HA mouse ascites 16B12 was used to visualize FH-tagged RIF1 as in A. Cell cycle stage was determined by DNA quantification (DAPI staining). E. Cell cycle distribution of the indicated cell lines, as determined by FACS quantification of EdU incorporation (S-phase) and DAPI staining (DNA amount). The average value of three independent clones per genotype is shown. Average of three experiments. Error bars indicate the standard error of the mean. *P* values are calculated using Wilcoxon test. F. Quantification by qRT-PCR of MERVL’s upregulation in the indicated genotypes, expressed as fold increase over MERVL levels in the same cell lines, before Cre induction. GAPDH was used to normalize. Average of two experiments is shown, each with 3 biological replicates for *Rif1-*Δ*PP1* and *Rif1-TgWT*, 2 for *Rif1-KO*, the parental cell line *Rif1-FH* and one reference for *Rif1-WT*. Error bars indicate standard deviations. Given the inherent clonal variability in the levels of MERVL expression, no quantitative conclusion is drawn from this experiment, except for presence/absence of the transcripts.

RIF1 deficiency in mESCs affects nuclear function at multiple levels. One of the features of Rif1-KO cells is the doubling of the population in G2, accompanied by a decreased S-phase population (Fig. 1E and ^13^). Our data show that *Rif1* hemizygosity (*Rif1-FH* and *Rif1-TgWT*) results in an altered cell cycle similar to RIF1 deficiency (Fig. 1E). Importantly, cell cycle-distribution in both *Rif1-*Δ*PP1* and *Rif1-TgWT* cell lines appears comparable to *Rif1-FH* cells. These results suggest that the defective cell cycle progression of *Rif1* null cells is not attributable to altered PP1 function but to insufficient levels of RIF1.

Loss of Rif1 function also results in the alteration of the gene expression profile in mESCs ^13^, including the de-repression of MERVLs ^38^, an effect that RIF1 shares with other epigenetic and DNA replication regulators ^53^. We therefore compared the level of MERVL RNA in *Rif1-WT*, *Rif1-KO*, *Rif1-TgWT, Rif1-FH* and *Rif1-*Δ*PP1* cells. After four days of deletion, MERVLs are upregulated not only in *Rif1-*Δ*PP1* and *Rif1-KO* cells, but, surprisingly, also in the hemizygous controls (*Rif1-TgWT* and *Rif1-FH* Fig. 1F), suggesting that, as for cell cycle progression, gene expression control is also sensitive to RIF1 dosage.

### RIF1-PP1 interaction is important for the replication-timing program

The most conserved function of RIF1 is the control of the replication-timing program and *Rif1-KO* cells show pronounced genome-wide changes in the temporal program of origin firing ^12,13^. As RIF1-PP1 interaction has been shown to be important, at least during the execution of the replication-timing program in S-phase, ^15,48–51^, expression of *Rif1*^Δ*PP1*^ should affect replication timing to a similar extent to *Rif1* deletion. In agreement with this prediction, hierarchical clustering of genome-wide replication timing shows that *Rif1-*Δ*PP1* and *Rif1-KO* mESCs cluster together, while *Rif1*^*+/+*^ (*Rif1-WT*) and control hemizygous (*Rif1-FH* and *Rif1-TgWT*) cells form a separate cluster (Fig. 2A and S2A).

**Figure 2.**
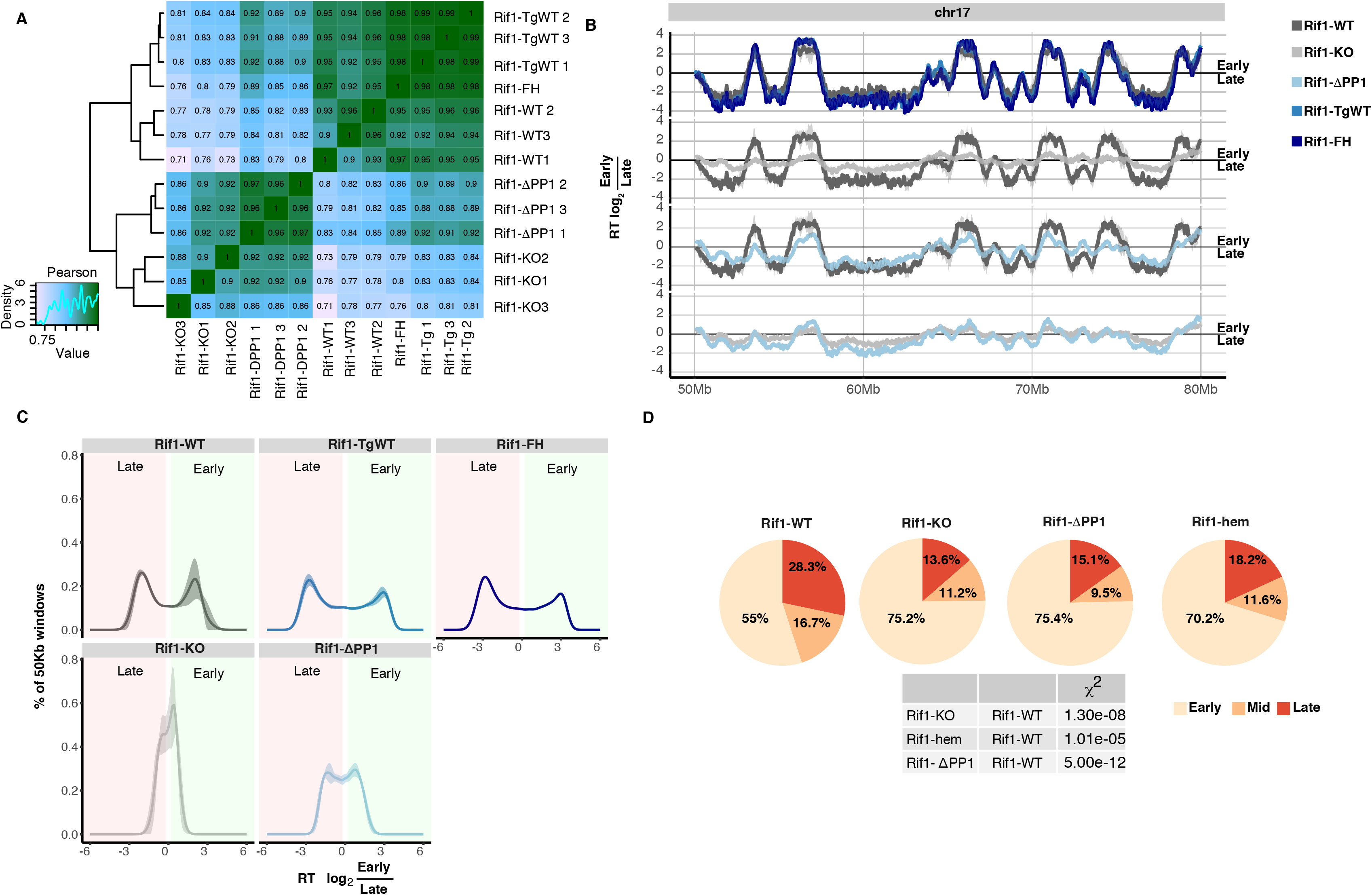
Effect of loss of RIF1-PP1 interaction on the replication-timing program and on the spatial distribution of replication foci. A. Hierarchical cluster analysis of Pearson correlation coefficient of genome-wide replication-timing (RT) profiles between replicas, bin size 50kb. The analysis shows preferential clustering of RT distribution from *Rif1-KO* and *Rif1-ΔPP1* lines, while RT distribution from *Rif1-WT* clusters with *Rif1-TgWT* and *Rif1-FH lines*. B. Representative RT profile from Chromosome 17. The solid line shows the average of three biological replicas, except for *Rif1-FH* (single, parental clone). RT scores are calculated as the log_2_ of the ratio between mapped reads in the early and late replicating fractions of the cell cycle over bins of 50Kb. C. Genome-wide distribution of 50kb genomic windows on the bases of their RT scores. Average of three independent lines per genotype is shown, except for *Rif1-FH*. Shaded areas represent standard deviations. RT scores from *Rif1-WT* and *Rif1* hemizygous lines (*Rif1-TgWT* and *Rif1-FH*) show a bimodal distribution, defining distinct early and late genomic regions. On the contrary, the distribution of RT scores from *Rif1-KO lines* shows a tendency towards a unimodal distribution, centered around zero. *Rif1-ΔPP1* lines display an increase of the windows with RT=0, but still a bimodal distribution of the RT values. D. The spatial distribution of replication foci (replication patterns) was visualized by EdU and DAPI staining. Cells were pulsed for 30 minutes with EdU and fixed. Examples in Fig. S4A. Pie charts show the relative distribution of S-phase cells (EdU positive) between replication patterns corresponding to early, mid and late S-phase. For each genotype, three independent lines and two separate experiments were blind-scored. As *Rif-FH* cells are a single cell line with no biological replicas (parental) and the results are very similar to the results from *Rif1-TgWT*, they were pooled (Rif1-hem). In the table, statistically significant differences are summarized. *P* values are calculated by χ^2^ test.

The replication profiles of both *Rif1-*Δ*PP1* and *Rif1-KO* mESCs appear similarly compressed around the zero (Fig. 2B and S2B), and a comparable fraction of the genome displays replication timing switches and changes (Fig. S3), suggesting an analogous loss of temporal control of origin firing in both cases. Importantly, the replication timing changes induced by the expression of *Rif1*^Δ*PP1*^ are not attributable to *Rif1* haploinsufficiency. In fact, the replication-timing profiles of *Rif1* hemizygous controls (*Rif1-FH* and *Rif1-TgWT*), are very similar to the wild type cells (*Rif1-WT*, Fig. 2B and C and Fig. S3, red boxes). Despite the similarities, however, the impact of loss of RIF1 versus loss of RIF1-PP1 interaction on the replication-timing program is qualitatively not identical. *Rif1*^Δ*PP1*^ expressing cells maintain a better degree of distinction between earlier and later replicating domains than *Rif-KO* (Fig. 2B, C and Fig. S2B). These data suggest that RIF1-dependent control of the replication-timing program could be not entirely exerted through PP1 and some other function of RIF1 partially could contributes as well.

### DNA replication timing is independent of the spatial distribution of replication foci

DNA replication takes place in a spatially organised manner ^54,55^, with the distribution of replication foci correlated to the time of replication ^56^. We have shown that in mouse primary embryonic fibroblasts (pMEFs), RIF1 deficiency induces changes of both the spatial distribution of replication foci and replication timing ^12^. We find a comparable effect in RIF1 deficient mESCs, with an increase of cells displaying an early-like replication pattern (Fig. 2D) despite there being no increase in the proportion of cells in early S-phase, as judged from the analysis of DNA content (Fig. S2C). In wild type cells, during early S-phase (defined as diffuse nucleoplasmic EdU and MCM3 staining and absence of histone H3 phosphorylated on Ser10-H3S10p), the replication pattern features many small replication foci throughout the nucleoplasm (Fig. S4A). In *Rif1-KO* cells, a similar distribution of replication foci (EdU) also appears aberrantly in cells in later S-phase (clusters of H3S10p signal, often at chromocenters, MCM3 discrete foci, larger-mid- or smaller-late and peripheral) (Fig. S4B). Normally at this stage, EdU signal appears as discrete foci of different sizes (Fig. S4A).

To study the effect of *Rif1* deficiency or expression of *Rif1*^Δ*PP1*^ on the total number of replication forks and their clustering, we have employed 3D-structure illumination microscopy (SIM-Fig. S5A). Since *Rif1* deletion and expression of *Rif1*^Δ*PP1*^ induce a loss of equivalence between replication foci distribution and replication timing, we could not analyse early, mid and late S-phase separately. *Rif1* deficiency results in an increase of the total number of replication forks (Fig. S5B) that could explain the apparent increase in the proportion of cells displaying early-like replication patterns. However, expression of *Rif1*^Δ*PP1*^ does not alter the total number of replication forks per nucleus (Fig. S5B), yet it causes an accumulation of early-like replication patterns that is comparable to *Rif1-KO* cells (Fig. 2D). Considering the different extent of the impact of loss of RIF1 (*Rif1-KO)* and loss of RIF1-PP1 interaction (*Rif1-*Δ*PP1*) on replication timing, this discrepancy in the effect on the total number of forks is interesting and indicates that the altered distribution of replication foci observed in both cell lines is not linked to the change of total number of replication forks. Moreover, by matching the total number of replication forks to the number of replication foci, we could not find a correlation between the number of forks per replication focus (Fig. S5C) and the changes of distribution of replication patterns (Fig. 2D). These data indicate that the increase of the proportion of cells with early-like replication patterns observed in *Rif1-KO* and *Rif1-*Δ*PP1* cells is not attributable to the de-clustering of the replication forks. Finally, our analysis shows that Rif1 hemizygosity (Rif1-hem=*Rif-FH*+*Rif1-TgWT*) has an impact on the spatial distribution of replication foci that is similar to, although milder, than *Rif1* deficiency or expression of *Rif1*^Δ*PP1*^ (Fig. 2D). However, hemizygosity does not result in an increased number of total replication forks or a measurable perturbation of the replication-timing program. These results suggest that spatial distribution of replication foci and the timing of replication can be uncoupled. In conclusion, loss or reduced RIF1 levels and loss of RIF1-PP1 interaction all impact on the distribution of replication foci, but not all affect replication timing.

### RIF1 dosage is important for nuclear compartmentalisation

The distribution of replication foci in the nucleus reflects the spatial organisation of the underlying chromatin. As a consequence, the altered spatial configuration of replication foci in *Rif1-hem* and *Rif1-*Δ*PP1* cells suggests that a reduced amount of RIF1 or loss of RIF1-PP1 interaction could affect chromatin organisation similarly to what we have shown for *Rif1* null cells ^13^, and irrespective of their effects on replication timing. We therefore analysed 3D chromatin organisation in *Rif1-KO*, *Rif1-*Δ*PP1* and *Rif1-hem* cells by Hi-C. We have previously shown by 4C that *Rif1* deficiency induces an increase of low-frequency contacts between TADs with different replication timing ^13^. In agreement with this, our Hi-C data indicate that *Rif1* deletion increases chromatin contacts *in cis,* especially at long range (>10Mbp) (Rif1-WT and Rif1-KO, Fig. 3A and B). The contacts gained preferentially involve late-replicating genomic regions associating with early-replicating regions (Fig. 4A) and RIF1-enriched regions gaining contacts with RIF1-poor genomic regions (Fig. 4B). These changes cannot be explained by the increased fraction of cells in G2 in *Rif1-KO* and *Rif1-*Δ*PP1* cells, as it was shown that chromosome compaction in G2/M favours the establishment of short-range interactions ^4^. Consequently, the increased proportion of the *Rif1-KO* and *Rif1-*Δ*PP1* cells in G2 will lead to an under-estimate of the true extent of the accumulation of long-range interactions. They suggest instead the alteration of the A/B compartmentalisation in mutant cells compared with *Rif1-WT* cells. Indeed, principle component analysis shows that *Rif1-WT* and *Rif1-KO* display a distinctly different compartment organisation (Fig. 4C). In agreement with previous data that have reported a more “open chromatin” ^12,13,17^, *Rif1*’s loss of function induces an expansion of the A compartment (and corresponding contraction of the B compartment, Fig. 4D) and compartment strength is weakened (Fig. 4E and F). Loss of RIF1-PP1 interaction induces alterations of chromatin organisation comparable to *Rif1-KO* (Fig. 4C), with a similar degree of expansion of the A compartment (Fig. 4D) and weakened compartment definition (Fig. 4E and F). This supports the conclusion that PP1 plays a key role in RIF1-dependent control of chromatin organisation. Unexpectedly, chromatin architecture in *Rif1* hemizygous cells (Rif1-FH and Rif1-TgWT) shows an intermediate but reproducible degree of change. Halving *Rif1* dosage, is sufficient to induce a gain of *in cis* contacts between distant genomic regions (Fig. 3A) of opposite replication timing (Fig. 4A). Accordingly, principal component analysis (PCA) of the overall A/B compartment organisation in *Rif1-TgWT* and *Rif1-FH* cells lies between with *Rif1* null and *Rif1-*Δ*PP1* cells on one side, and *Rif1-WT* (Fig. 4C) on the other, with an expansion of the A compartment (Fig. 4D) and weakened compartmentalisation (Fig. 4E and F) that is intermediate between *Rif1-WT* and *Rif1-KO/Rif1-*^Δ*PP1*^ cells. These data indicate that chromatin architecture is exquisitely sensitive to RIF1 dosage, and that decreasing the levels of RIF1 induces a progressive alteration of nuclear organisation. This is in striking contrast with the effect of varying RIF1 levels on the regulation of replication timing.

**Figure 3.**
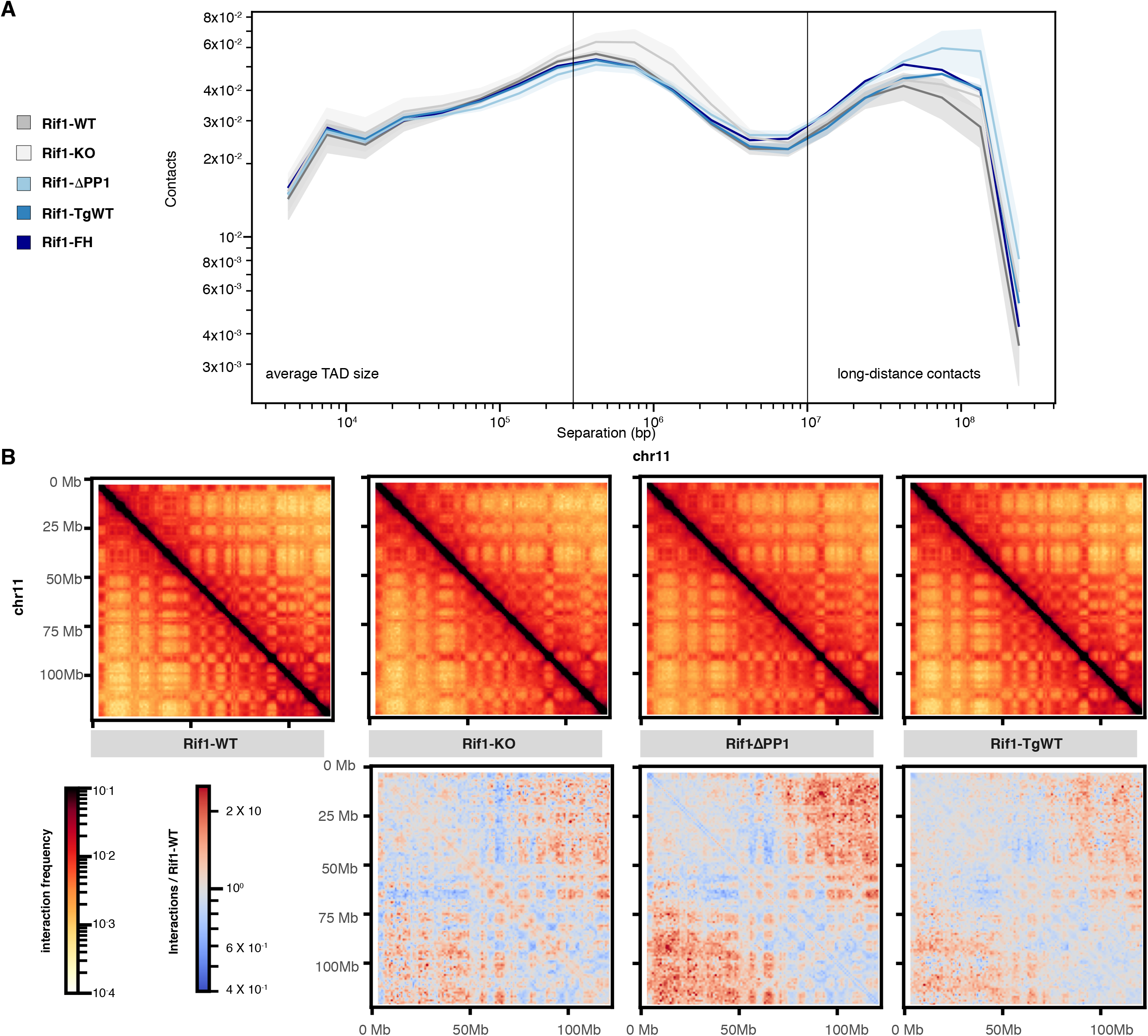
RIF1 spatially confines chromatin contacts in a dose-dependent manner. A. Normalised contact frequency versus genomic distances for Hi-C reads normalised. Three biological replicas per genotype, except for *Rif1-FH*, are shown. *Intra*-TADs contacts (line at a median TAD’s size of approximately 0.3Mbp), and long-range (over 10Mbp apart) are indicated. Shaded areas represent standard deviations. B. Representative distribution of the median number of *in cis* chromatin contacts per indicated position (arbitrary units) within the specified region of Chromosome 11. Three independent clones per genotype were used. Upper: log(balanced HiC signals). Lower: log((balanced HiC signals (indicated line/*Rif1-WT*)). Red indicates a gain of interactions over *Rif1-WT*, while blue represents a loss.

**Figure 4.**
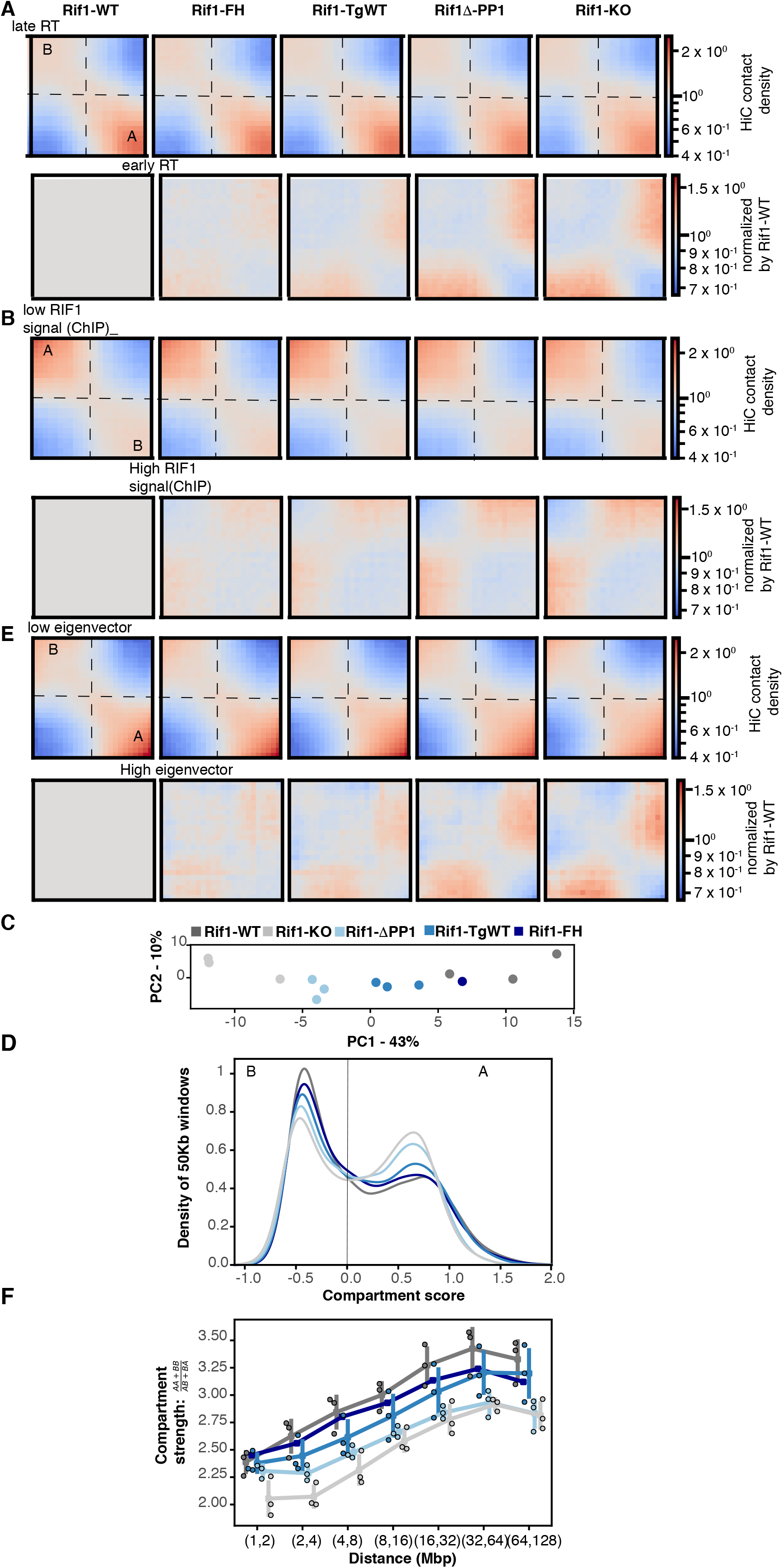
Segregation of A and B nuclear compartments is sensitive to Rif1 dosage. A. Top row: Saddle plot of Hi-C data, binned at 250 kb resolution for loci ranked by their replication timing. An increase in contacts of genomic positions of opposite replication timings is progressively more evident from *Rif1-WT* to *Rif1-KO*. The triplicates for each genotype (except for *Rif1-FH*) were combined. Lower row: Hi-C for each genotype, data normalized to *Rif1-WT*. Red indicates a gain in contacts. A and B indicates the compartments. B. Top row: Saddle plot of Hi-C data, binned at 250 kb resolution for loci ranked by their association with RIF1 ^13^. An increase in contacts between RIF1-associated and RIF1-devoided genomic positions is progressively more evident from *Rif1-WT* to *Rif1-KO*. The triplicates for each genotype (except for *Rif1-FH*) were combined. Lower row: Hi-C for each genotype, data normalized to *Rif1-WT*. Red indicates a gain in contacts. A and B indicates the compartments. C. Principle component analysis of A/B compartmentalisation for the indicated genotypes in triplicate, except for *Rif1-FH.* D. Distribution of genomic regions of 250kb windows between the A and B compartment. Average of three biological replicates is shown, except for the parental line *Rif1-FH*. E. Top row: Saddle plot of Hi-C data, binned at 250 kb resolution for loci ranked by their eigenvector values. An increase in contacts between genomic positions in different compartments is progressively more evident from *Rif1-WT* to *Rif1-KO*. The triplicates for each genotype (except for *Rif1-FH*) were combined. Lower row: Hi-C for each genotype, data normalized by *Rif1-WT*. Red indicates a gain in contacts. A and B indicates the compartments. F. Compartment strength variation with distance for the indicated genotypes. Individual values for the three biological replicates are represented by the outlined circles, except for *Rif1-FH*. The bars represent standard deviations.

## Discussion

The remarkable coincidence of spatial distribution and replication timing of different portions of the genome, at multiple levels of organisation and throughout evolution, has encouraged the idea of a causal relationship between nuclear architecture and replication timing. At a molecular level, their covariation – for example during cell fate determination and embryonic development – finds a confirmation in their co-dependence on RIF1. In this work, we show that both aspects of nuclear function depend upon the interaction between RIF1 and PP1. However, 3D organisation of chromatin contacts and replication timing show a different degree of dependency on RIF1-PP1 interaction and are differentially influenced by RIF1’s dosage. The loss of RIF1-PP1 interaction affects the compartmentalisation of chromatin contacts comparably to a complete loss of RIF1 function, while it only partially recapitulates the effects of *Rif1*^*−/−*^ on the control of replication timing. In addition, the former is sensitive to RIF1’s dosage, while *Rif1’s* haploinsufficiency does not affect the latter. In summary, there is a clear distinction in two groups: 1. replication timing is only affected by complete loss of RIF1. 2. On the contrary, nuclear compartmentalisation, long-range chromatin contacts, replication foci spatial organisation and MERVL repression all show sensitivity to RIF1 dosage, with the effect of lack of RIF1-PP1 interaction worsening the effect of hemizygosis clearly for the long-range chromatin interactions and nuclear compartmentalisation. We hypothesise that the reason for the difference of the effect of RIF1-PP1 loss of interaction between chromatin contacts/nuclear compartmentalisation and replication foci distribution/MERVL overexpression, is that chromatin contacts/nuclear compartmentalisation are the primary features affected by loss of RIF1-dependent dephosphorylation of critical substrate/s, that directly or indirectly control them. The changes of replication foci distribution are a more indirect way to visualise the same changes (therefore less sensitive), and changes in gene expression (MERVL upregulation, in this case) could also be an indirect consequence of these architectural changes, as we had already hypothesised ^12,13^. We have indeed shown that the transcriptome is only altered after few cell cycles in absence of RIF1, while nuclear architecture changes are an immediate consequence of RIF1 absence, in the first cell cycle after *Rif1* deletion^13^.

RIF1 is known to multimerise ^45,47,57^ and to interact with the nuclear lamina ^13,42^. RIF1 multimers could act as a sub-stochiometric platform, interacting with different regulators of replication timing, in addition to PP1. In this case, the consequences of the complete loss of RIF1 function on the replication-timing program would amount to the sum of perturbation of multiple pathways that control the timing of origin activation. For example, RIF1-ΔPP1 may only specifically interfere with the PP1-dependent control of DDK (Dbf4-dependent kinases) activity at origins, while other RIF1 interactors may contribute to the epigenetic control of origin activation. The proteins associated with RIF1 are enriched for chromatin and epigenetic regulators ^52^, and the contribution of histone modifiers to the control of replication timing has long been recognised ^58–63^. However, an understanding of the effect of *Rif1* deletion on the epigenetic landscape is still missing, leaving this hypothesis currently hard to test ^12,13,38,64^. In the context of chromatin architecture organisation, RIF1 multimers could directly participate in the creation of local scaffolds that restrict chromatin mobility or, alternatively, could regulate other proteins with this role. In either case, a reduction of RIF1 dosage could have structural, quantitative consequences.

Our results identify RIF1 as a molecular link, a point of convergence and co-regulation. We propose that RIF1, specifically, and not generic nuclear architecture, coordinates the replication-timing program with nuclear 3D organisation. In agreement with this view, recent data show that cohesins and CTCF are not involved in the regulation of replication timing ^7,65^ and that the definition of A/B compartments and Early/Late replicating domains is uncoupled at the time of zygotic genome activation in zebrafish ^66^. Altogether, these data suggest that replication timing and nuclear architecture, or at least 3D organisation of chromatin contacts and spatial distribution of replication foci, are not linked by a causative relationship. Yet, they are coregulated, both during cell cycle and embryonic development, and RIF1 is a point of convergence. Having established this, is an important step to start addressing the fundamental question of why this coordination is important. During embryonic development in different organisms, for example in *Drosophila melanogaster*, replication timing ^16^ and TADs definition both emerge around the time when zygotic transcription starts ^67^. Could uncoupling these two events have consequences on gene expression? We can alter chromatin organisation, leaving replication timing intact, by halving RIF1 dosage. This affects cell cycle progression and the repression of MERVLs (this work). In a complementary approach, it has been shown that alteration of replication timing by overexpression of limiting replication factors during early *Xenopus laevis* development, that presumably leaves nuclear architecture intact, affects the onset of zygotic transcription and the transition into gastrulation ^68^. It is therefore tempting to speculate that the covariation of replication timing and nuclear architecture could be important to coordinate gene expression and the choice of origins of replication.

## Methods

### Mouse embryonic stem cell derivation

Mouse embryonic stem cell (ESC) cells were derived as described in (Foti, Gnan et al. 2016), with the addition of 1 μM MEK inhibitor PD0325901 and 3 μM GSK3 inhibitor CHIR99021 (MRC PPU Reagents and Services, School of Life Sciences, The University of Dundee) in the culture media, from the start of the protocol. Rif1^FH/flox^ Rosa26^Cre-ERT/+^ ESCs were derived by crossing Rif1^flox/+^ Rosa26^Cre-ERT/Cre-^ERT ^19^ with Rif1^FH/FH 12^ mice. The Rif1^FH^ allele was specifically targeted in the parental line Rif1^FH/flox^ Rosa26^Cre-ERT/+^ (Rif1-FH). Integrants were selected by hygromycin resistance. The targeting vector encodes for a codon-optimized cDNA of RIF1 (exon 8 to exon 36). Hygromycin-resistant colonies were screened for correct targeting of the Rif1^FH^ allele by Southern blot (EcoRV digest) and using a PCR-amplified probe (primers in Table 1).

### Cell manipulation

ESCs were grown at 37°C in 7.5 % CO_2_ in Knockout DMEM (Gibco 10829-018), containing 12.5% heat-inactivated fetal bovine serum (Pan-Biotech), 1 % non-essential amino acids (Gibco 11140-035), 1% Penicillin/Streptomycin (Gibco 15070063), 0.1 mM 2-Mercaptoethanol (Gibco 31350-010), 1% L-Glutamine (Gibco 25030024)), supplemented with 1 μM PD0325901 and 3 μM CHIR99021 and 20 ng/ml leukemia inhibitory factor (LIF, EMBL Protein Expression and Purification core facility).

Experiments were carried out each time from a frozen vial of cells, at least two passages after thawing. 5.2·10^6^ cells for Rif1-WT and 6.5·10^6^ for Rif1-KO lines, per 15 cm plate (or the equivalent for different sized plates) were plated at day zero, when treatment with 200 nM 4-hydroxytamoxifen (OHT, Sigma H7904) started. Fresh medium with OHT was added after 48 hours. Cells were collected about 96 hours after starting OHT treatment.

### Replication Timing analysis

Cells were pulsed for 2 hours with 10 μM BrdU, collected and fixed in 70 % ethanol. Processing was as described in ^69^. Fastq files were aligned using Bowtie2 version 2.2.6 on mm10 as a reference genome. SAM files were converted into BAM files and sorted using Samtools version: 1.3.1. bamCompare version 3.1.3 was used to create bedgraph files with 50 kb and 1 kb binning of the log_2_ ratio of the early and late fraction. Duplicated reads were excluded from the computation of the bedgraph files as well as reads mapped on XY chromosomes. The two fractions were normalized as reads per millions (RPM). Plots and data manipulation were carried out R version 3.5.1. The original names of the cell lines used in these experiments, included in the name of the Repli-seq raw files are: RFHF14 = Rif1-FH, 14 tgWT A7 = Rif1-TgWT 1, 14 tgWT H4 = Rif1-TgWT 2, 14 tgwt H6 = Rif1-TgWT 3, 14 ΔP G11 = Rif1-ΔPP1 1, 14 ΔP H1 = Rif1-ΔPP1 2, 14 ΔP H2 = Rif1-ΔPP1 3, ESC B = Rif1-WT 1, ESC F = Rif1-WT 2, ESC H = Rif1-WT 3, ESC 5 = Rif1-KO 1, ESC 18 = Rif1-KO 2, ESC 24 = Rif1-KO 3.

### Cell cycle distribution analysis

After four days of OHT treatment, cells were pulsed for 30 minutes with 10 μM EdU (Invitrogen A10044). Cells were then washed with cold DPBS (Thermo Fisher Scientific 14190094), collected, counted and fixed in 75 %. EtOH Samples were kept at −20 °C for at least overnight. 7.5·10^5^ cells were then processed for click-chemistry detection of EdU. After washing in cold DPBS, cells were permeabilised in DPBS/1% FBS/0.01 % Triton X-100 (Sigma93426-250ML) for 10 minutes on ice. After washing twice, cells were incubated in 900 μl of DPBS with 10 mM Na-Ascorbate (Sigma A7631-25G), 1 μM Alexa Fluor 647 Azide (Thermo Fisher Scientific A10277) and CuSO_4_ 0.1 M (Sigma C1297) for 30 minutes at room temperature in the dark, rotating. Cells were washed in DPBS/1%FBS/0.5% Tween 20 (Sigma P9416-100ML) for 10 minutes and then twice in cold DPBS/1% FBS. After 1 hour incubation in 300 μl of DPBS/1%FBS /DAPI 2.5 g/ml (Thermo Fisher Scientific D1306), the samples were analyzed using an LSR II FACS (BD). The data acquired were analysed using Flowjo software and plotted in R 3.5.1. To calculate the percentages of cells in early, mid and late S-phase in Fig. S2C, we have defined the S-phase substages based on the intensities of the PI/EdU signals in the wild type, drawn the gates and applied them to all the samples ^12^.

### Intra-cellular FACS staining for ^HA^Rif1

After four days of OHT treatment, cells were collected and counted. 3·10^6^ cells were fixed in 400 μl of DPBS/2% Paraformaldehyde (Sigma P-6148) for 10 minutes at room temperature shaking. Paraformaldehyde was then diluted to 0.2% and next cells were washed in cold DPBS. After 2 minutes permeabilisation in 200 μl PBS-Triton X-100 0.1%, cells were incubated 5 minutes in saponin solution (COMPONENT E from kit C10424, Thermo Fisher Scientific) at room temperature and anti-HA antibody (Covance monoclonal HA.11 clone 16B12 #MMS-101R, RRID:AB_291262) was added at 1:500. After 1 hour at room temperature rotating, cells were washed twice in DPBS/2% FBS, resuspended in 200 μl of saponin solution with goat anti-mouse Alexa Fluor 647 1:1000 (Thermo Fisher Scientific A-21235, RRID:AB_2535804) and incubated for 1 hour rotating in the dark. After washing twice samples were resuspended in 400 μl of saponin solution with DAPI 2.5 g/ml (Thermo Fisher Scientific D1306) and analyzed on an LSR II FACS (BD). Data were processed using R version 3.5.1. The confidence intervals (CI) of the median shown in Fig. 1B were manually calculated by bootstrap.

For the FACS analysis of RIF1’s chromatin association, the samples were processed as above, except, fixation was preceded by 3 minutes incubation in CSK buffer (25 mM HEPES pH 7.4, 50 mM NaCl, 1 mM EDTA, 3 mM MgCl2, 300 mM sucrose, 0.5% Triton X-100 and complete protease inhibitor cocktail tablet). Pre-extracted cells were subsequently fixed in 3%PFA/sucrose for 30 minutes at room temperature shaking.

## Acknowledgments

We would like to acknowledge Martin Waterfall from the IIIR Flow Cytometry Core Facility, University of Edinburgh; David Kelly from the COIL facility, WTCCB, University of Edinburgh; Vladimir Benes and the Genomic Core Facility at EMBL Heidelberg; Philip Hublitz from Gene expression Facility, EMBL Monterotondo; Violetta Parimbeni for mouse husbandry, EMBL Monterotondo. SG was funded by ERC consolidator award 726130 to SCBB; EC was supported by the Erasmus Program. EE received funding from the European Union’s Horizon 2020 research and the Marie Skłodowska-Curie Individual Fellowship grant agreement No. 660985 and from the ERC consolidator award 726130 to SCBB. IMF was funded by the Darwin Trust of Edinburgh. SCBB thanks the ERC, DMG thanks NIH grant GM083337, WAB is funded by a Medical Research Council University Unit programme grant [MC_ UU_00007/2]. MCC was funded by the Deutsche Forschungsgemeinschaft (DFG, German Research Foundation) – Project-ID 393547839 – SFB 1361 TP06 and DFG grant CA 198/9-2.

## Author Contributions

SG has created the cellular system, performed the majority of the experiments and the bioinformatics analysis. IMF has helped with HiC experiments and analysis. KNK has performed the replication timing measures. EC and NC have stained and scored with SG replication-timing patterns. PW, AR and AM have respectively stained cells, acquired 3D-SIM images and performed the analysis. EE has analysed MERVL expression. WAB, DMG, MCC have supported the work with personnel, resources, scientific discussions and critical reading of the manuscript. SCBB has conceived the project, performed some of the experiments and written the manuscript.

