## supplemental info for "Nuclear organisation and replication timing are coupled through RIF1-PP1 interaction"

**Table of contents of Supplemental Material**

Supplemental Figure Legend
Supplemental Figures
Supplemental Methods

**Supplemental Information**

**Fig. S1 Generation of ESCs expressing RIF1-ΔPP1.** Related to Fig. 1.

A. In Rif1$^{FH/flox}$ ESCs, the Rif1$^{FH}$ allele was targeted. The targeting construct allows knocking-in Rif1's codon-optimised cDNA (exons 8 to 36), either wild type or carrying the SAAA/RVSF (Rif1$^{\Delta PP1}$) mutations of the SILK and RVSF motifs, residues 2128–2131 and 2150–2153. The homology arms included in the construct to target the *Rif1* locus are shaded in grey. The targeted alleles are indicated in the figure as $^{FH}$Rif1$^{TgWT}$ or $^{FH}$Rif1$^{\Delta PP1}$ respectively. Splicing between genomic-encoded exon 7 and cDNA-encoded exon 8 allows expression of Rif1-TgWT or Rif1-ΔPP1 respectively. Restriction sites and probe used to identify by Southern blot the correct insertion of the targeting construct in Rif1$^{FH}$ allele and not the Rif1$^{flox}$ allele are indicated. The map of Rif1$^{flox}$ allele indicating the expected sizes after restriction digest and Southern blot analysis is shown.
B. Southern blot confirmation of correct integration of the targeting constructs in the cell lines employed in this work.
C. Western blot analysis of RIF1 levels after four days of hydroxytamoxifen treatment, to induce Cre-mediated deletion of Rif1$^{flox}$ allele. Proteins were extracted from untagged *Rif1-WT* (Rif1$^{+/+}$); *Rif1-KO* (Rif1$^{flox/flox}$); FLAG-HA2(FH) knock-in tagged Rif1 hemizygous: *Rif1-FH* (Rif1$^{FH/flox}$); FH-tagged targeted *Rif1-TgWT* hemizygous (Rif1$^{TgWT/flox}$); FH- tagged targeted *Rif1-ΔPP1* hemizygous (Rif1$^{\Delta PP1/flox}$). Anti-mouse Rif1 polyclonal rabbit antibody 1240 (anti-RIF1) was used to detect both FH-tagged and untagged proteins. Mouse ascites 16B12 (anti-HA) detects only FH-tagged RIF1. SMC1 levels were used as loading control.
D. Independent experiment as in Fig. 1D. Quantitative analysis of the levels of chromatin-associated FH-tagged RIF1 throughout the cell cycle (one representative experiment), measured by FACS staining. Cytoplasmic and nucleoplasmic proteins were pre-extracted before fixing chromatin-associated proteins. Anti-HA mouse ascites 16B12 was used to visualize FH-tagged RIF1 as in A. Cell cycle stage was determined by DNA quantification (DAPI staining).

**Fig. S2 Replication timing is affected by lack of RIF1 or of RIF1-PP1 interaction, not RIF1 dosage.** Related to Fig. 2.

A. Principle component analysis of RT for the indicated genotypes in triplicate, except for RIF1-FH.
B. Representative RT profile from one line per genotype at 1kb resolution.

C. Relative distribution of S-phase cells (EdU positive) between DNA contents corresponding to early, mid and late replication, as determined by DAPI quantitation (FACS). The average value of three independent clones with the same, indicated genotypes is shown. Average of three experiments. Error bars indicate standard error of the mean. *P* values are calculated using Kruskal–Wallis test.

**Fig. S3 RIF1-ΔPP1's impact on the replication-timing program does not entirely recapitulate the consequences of RIF1 loss of function.** Related to Fig. 2.

Percentage of 50kb-windows changing RT from Early-to-Late S-phase (EtoL, from a positive to a negative RT value), Early-to-Earlier (to Earlier, thus more positive value value of RT), from Late-to-Early (LtoE, from a negative to a positive RT value), and from Late-to-Later S-phase (to Later, thus more negative value of RT), over increasing thresholds. The black vertical lines indicate ΔRT=1, the threshold delimiting significative RT changes (above the experimental and clonal variation; see for example (*Rif1-FH*)-(*Rif1-TgWT*) and [13]). The red boxes highlight the comparison between wild type and hemizygous lines: all the differences are ΔRT<1.

**Fig. S4 Classification of dynamic spatial distribution of replication foci in mESCs.** Related to Fig. 2.

A. G1, G2 and S-phase substages classification. The different stages are distinguished by the combination of EdU staining (grey), anti-MCM3 (red) and anti-histone H3 phosphorylated on Ser10 (H3S10p, green) immunofluorescence. G1 cells display a diffuse nucleoplasmic MCM3 signal, but no EdU or H3S10p signal. Early S-phase is characterised by diffuse nucleoplasmic MCM3 and EdU staining, with no H3S10p signal. In mid S-phase MCM3 and EdU signal concentrate more in the chromocenters and H3S10p starts appearing. In late S-phase there are peripheral or sparse MCM3 and EdU signals, with H3S10p signal becoming very prominent, especially at chromocenters.
B. Examples of aberrant replication patterns in *Rif1-KO* cells. H3S10p signal, together with MCM3 located at chromocenters, normally associated with mid-late S-phase EdU patterns, is in these examples associated with diffuse nucleoplasmic EdU patterns resembling early S-phase.

**Fig. S5 Analysis of the number and clustering of replication forks in mESCs.** Related to Fig. 2.

A. Example of image processing and workflow to evaluate the number of replication forks (nano-replication foci, nano-RFi) versus the number of fork clusters ((pseudo)-wide-field replication focus, pWF-RFi). pWF-Rfi contain usually more than one nano-RFi, but there is a fraction of nano-RFi not associated with a pWF-RFi (yellow arrowheads).

B. Number of replication forks per cell, for two independent biological replicates per genotype (except for *Rif1-WT*, where only one clone was analysed). The line represents the median values and the 95% confidence intervals. *P* value is calculated using Kruskal Wallis test. The forks were visualised by anti-BrdU staining and 3D-SIM image acquisition.

C. Boxplots showing the distribution of the number of replication forks per replication focus. *P* values are calculated using Kruskal Wallis test. (0.05> *P* value >0.01 == *, 0.01 >= *P* value >0.001 == **, 0.001 >= *P* value >0.0001 == ***, *P* value < 0.0001 == ****)

### Figure S1

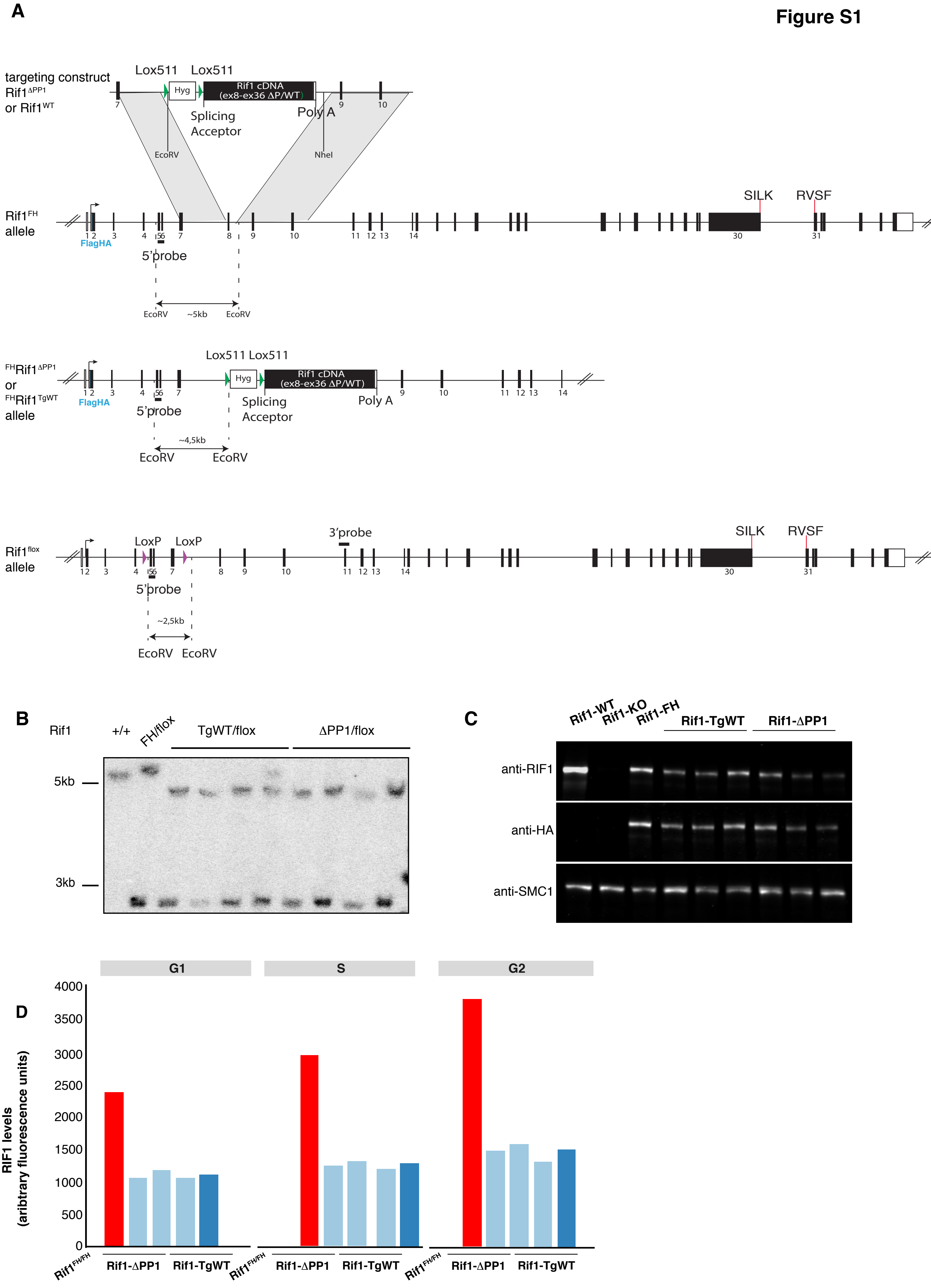
A
targeting construct
Rif1 ΔPP1
or Rif1 WT
Lox511
Lox511
Hyg
Rif1 cDNA
(ex8-ex36 ΔP/WT)
Splicing
Acceptor
Poly A
EcoRV
NheI
Rif1 FH
allele
FlagHA
5'probe
EcoRV
~5kb
EcoRV
SILK
RVSF
FH Rif1 ΔPP1
or
FH Rif1 TgWT
allele
~4,5kb
Rif1 flox
allele
LoxP
LoxP
3'probe
~2,5kb
B
Rif1
+/+
FH/flox
TgWT/flox
ΔPP1/flox
5kb
3kb
C
Rif1-WT
Rif1-KO
Rif1-FH
Rif1-TgWT
Rif1-ΔPP1
anti-RIF1
anti-HA
anti-SMC1
D
G1
S
G2
RIF1 levels
(aribtrary fluorescence units)
4000
3500
3000
2500
2000
1500
1000
500
0
Rif1 FH/FH
Rif1-ΔPP1
Rif1-TgWT

**Figure S2**

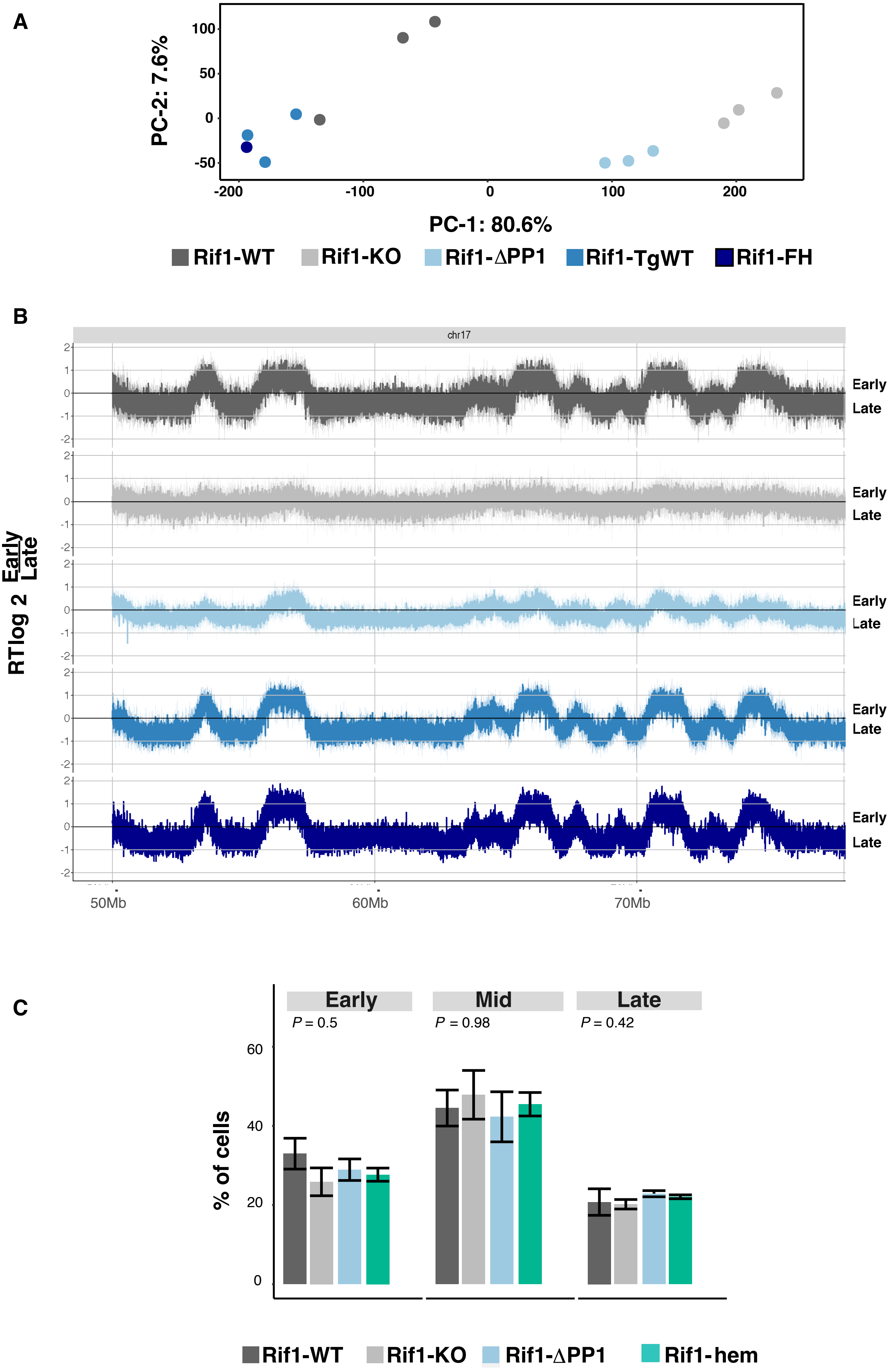
A
PC-2: 7.6%
100
50
0
-50
-200
-100
0
100
200
PC-1: 80.6%
Rif1-WT
Rif1-KO
Rif1-ΔPP1
Rif1-TgWT
Rif1-FH
B
chr17
RT log 2 Early/Late
Early
Late
50Mb
60Mb
70Mb
C
Early
Mid
Late
P = 0.5
P = 0.98
P = 0.42
% of cells
60
40
20
0
Rif1-WT
Rif1-KO
Rif1-ΔPP1
Rif1-hem

**Figure S3**

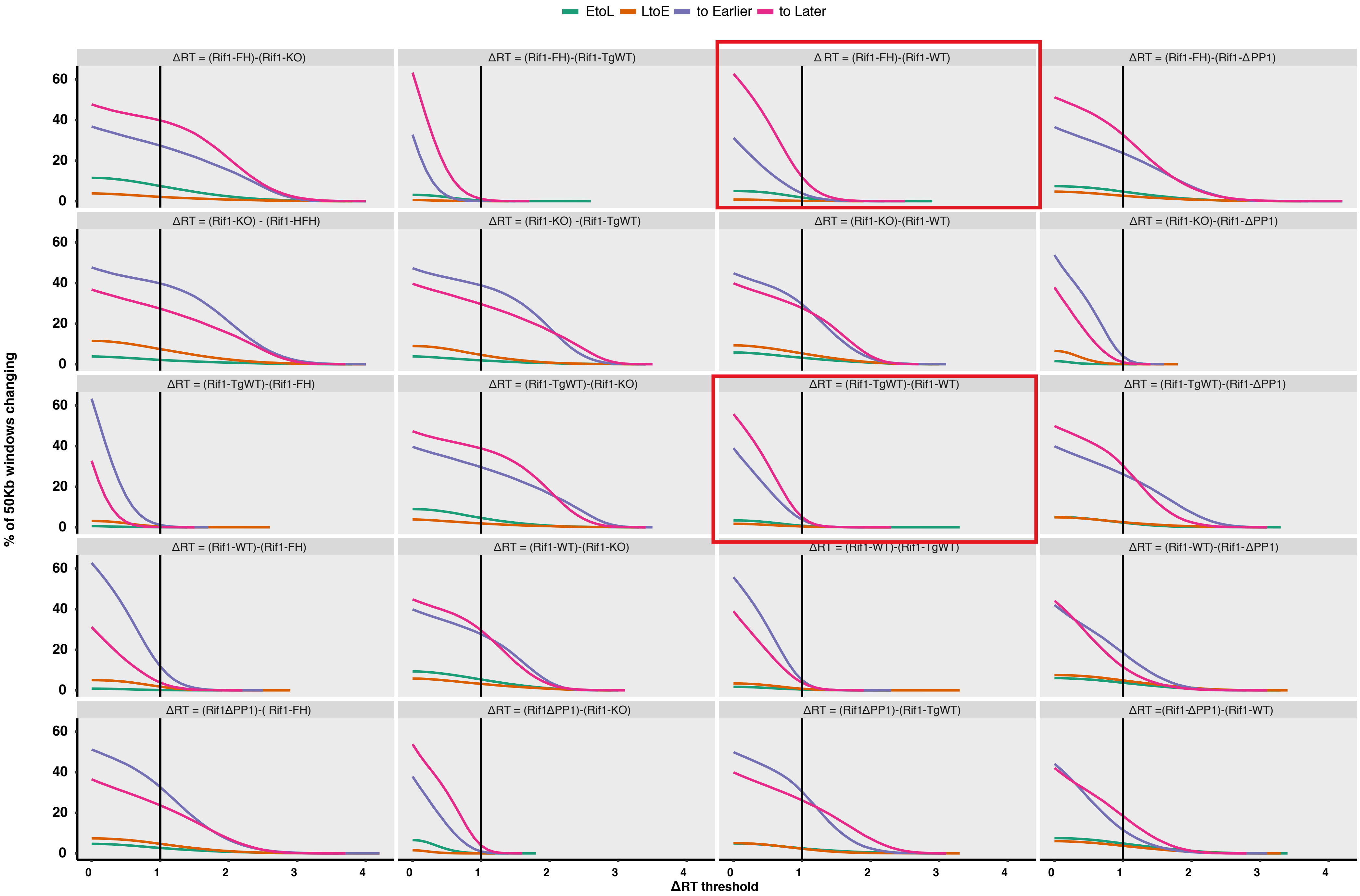

EtoL
LtoE
to Earlier
to Later
ΔRT = (Rif1-FH)-(Rif1-KO)
ΔRT = (Rif1-FH)-(Rif1-TgWT)
Δ RT = (Rif1-FH)-(Rif1-WT)
ΔRT = (Rif1-FH)-(Rif1-ΔPP1)
ΔRT = (Rif1-KO) - (Rif1-HFH)
ΔRT = (Rif1-KO) -(Rif1-TgWT)
ΔRT = (Rif1-KO)-(Rif1-WT)
ΔRT = (Rif1-KO)-(Rif1-ΔPP1)
ΔRT = (Rif1-TgWT)-(Rif1-FH)
ΔRT = (Rif1-TgWT)-(Rif1-KO)
ΔRT = (Rif1-TgWT)-(Rif1-WT)
ΔRT = (Rif1-TgWT)-(Rif1-ΔPP1)
ΔRT = (Rif1-WT)-(Rif1-FH)
ΔRT = (Rif1-WT)-(Rif1-KO)
ΔRT = (Rif1-WT)-(Rif1-TgWT)
ΔRT = (Rif1-WT)-(Rif1-ΔPP1)
ΔRT = (Rif1ΔPP1)-( Rif1-FH)
ΔRT = (Rif1ΔPP1)-(Rif1-KO)
ΔRT = (Rif1ΔPP1)-(Rif1-TgWT)
ΔRT =(Rif1-ΔPP1)-(Rif1-WT)
% of 50Kb windows changing
0
20
40
60
0
1
2
3
4
ΔRT threshold

Figure S4

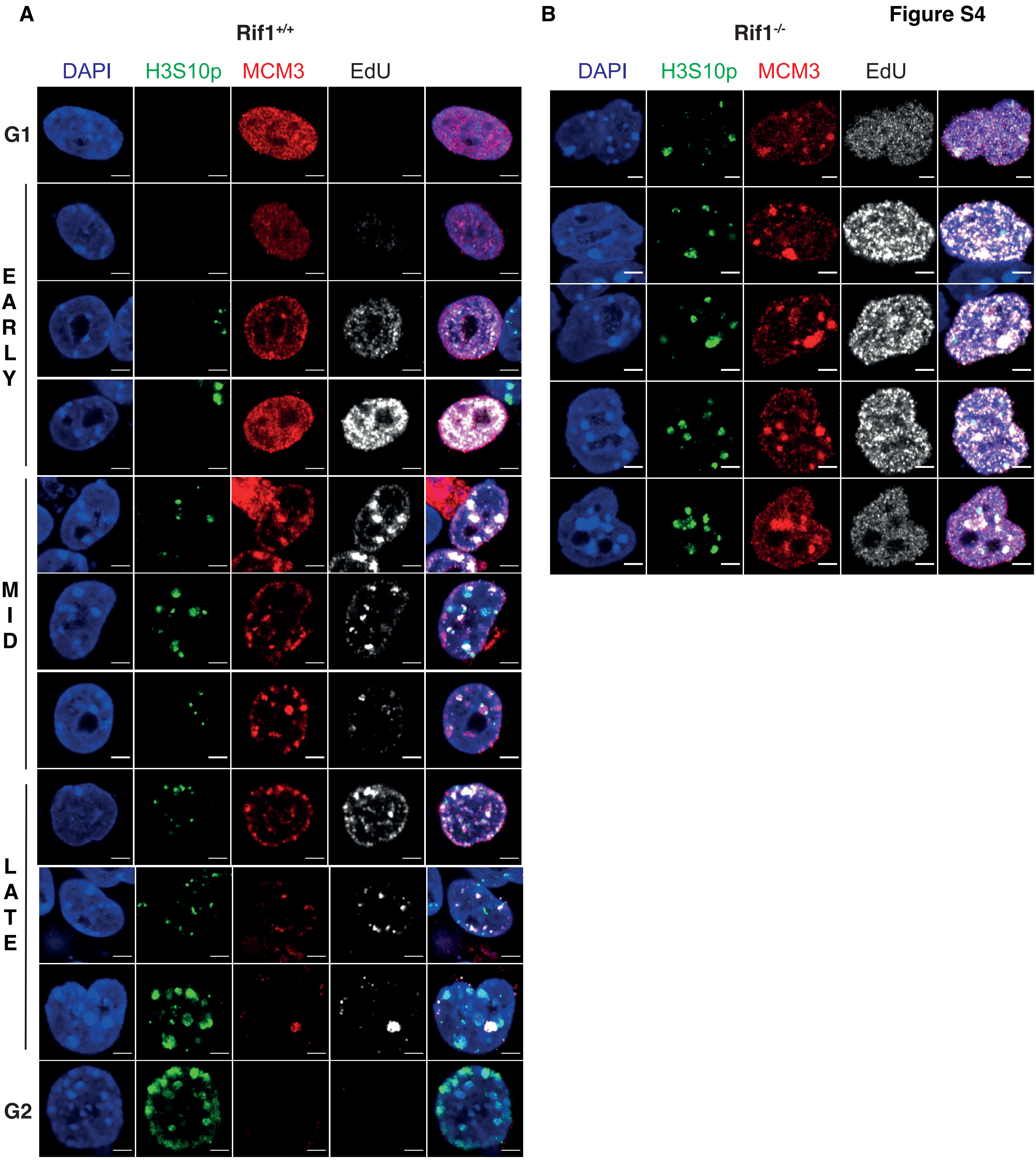
A
Rif1+/+
DAPI
H3S10p
MCM3
EdU
G1
EARLY
MID
LATE
G2
B
Rif1-/-
DAPI
H3S10p
MCM3
EdU

**Figure S5**

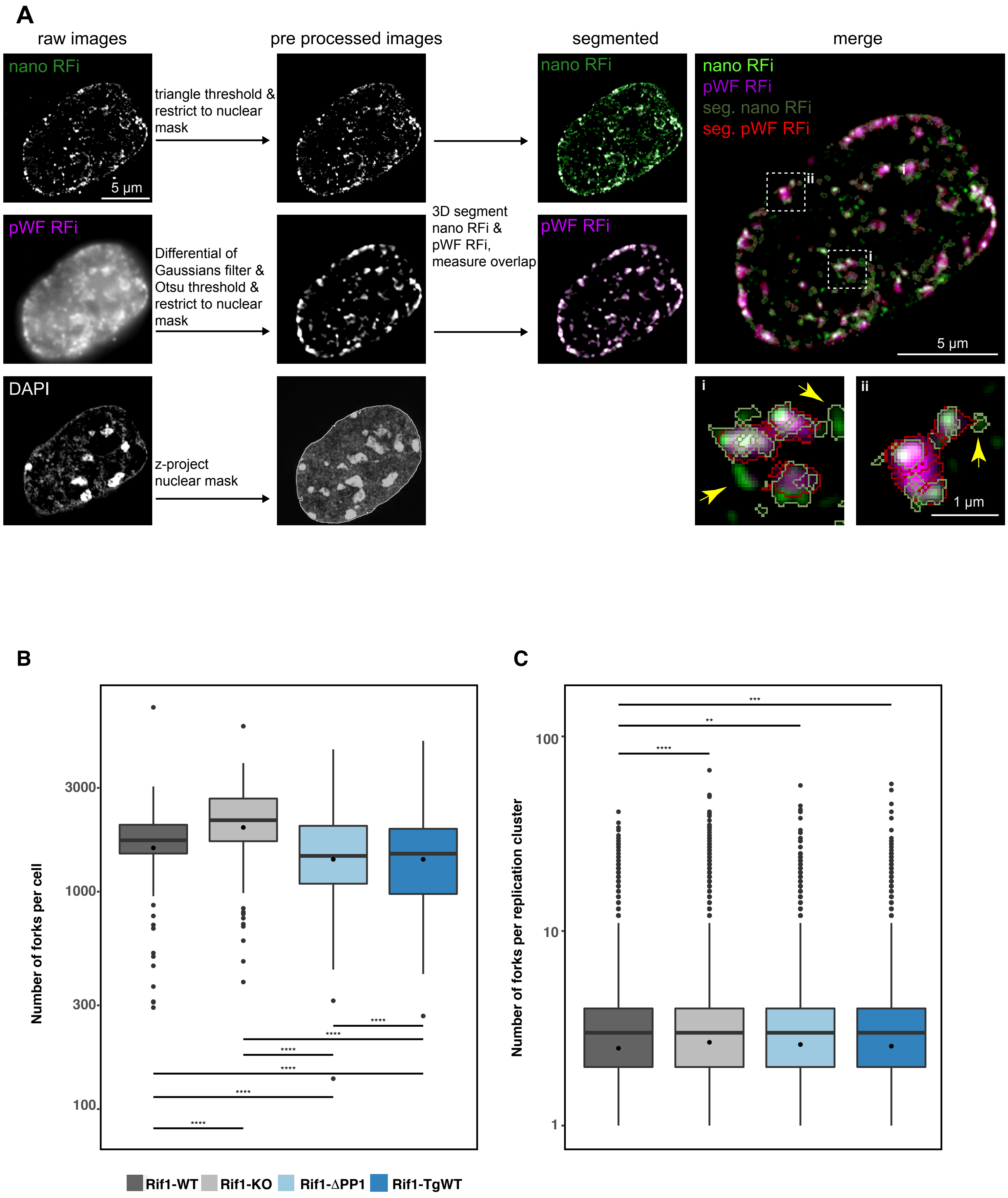
A
raw images
pre processed images
segmented
merge
nano RFi
5 µm
triangle threshold & restrict to nuclear mask
pWF RFi
Differential of Gaussians filter & Otsu threshold & restrict to nuclear mask
DAPI
z-project nuclear mask
3D segment nano RFi & pWF RFi, measure overlap
nano RFi
pWF RFi
seg. nano RFi
seg. pWF RFi
i
ii
5 µm
1 µm
B
Number of forks per cell
3000
1000
300
100
****
C
Number of forks per replication cluster
100
10
1
****
**
***
Rif1-WT
Rif1-KO
Rif1-ΔPP1
Rif1-TgWT

**Supplemental Methods**

**Hi-C**

After four days of OHT treatment, cells were collected and counted. Cells were washed twice in cold DPBS and resuspended in full media. Samples were crosslinked for 10 minutes rotating at room temperature in 1% formaldehyde ($10^6$ cells/ml). Crosslinking was stopped by adding glycine at the final concentration of 0.2 M for 5 minutes at RT. Samples were washed twice in cold DPBS and pellets were snap frozen. 2-5 × $10^6$ cells per sample were lysed in 10 mM Tris-HCl pH 8.0, 10 mM NaCl, 0.2% IGEPAL (Sigma, I3021) supplemented with protease inhibitor (Thermo Scientific, 78430) for 15 minutes on ice. Nuclei were washed twice in cold lysis buffer, resuspended in 50 µl of NEBuffer 3.1 (NEB, B7003S) supplemented with 0.3% SDS and incubated at 62°C for 10 minutes. After diluting and quenching the SDS by adding 57.5 µl of NEBuffer 3 and 12.5 µl of 20% Triton X-100 (Sigma, 93443), samples were incubated at 37°C for 60 minutes. Nuclei were spun and incubated in 250 µl of 1× DpnII buffer with 600 U of DpnII restriction enzyme (NEB, R0543L) overnight at 37°C. An additional 200 U of DpnII were added to each sample the following day and incubated for a further 2 hours.

After heat inactivation of DpnII for 20 min at 65°C, restriction fragments ends were filled by DNA polymerase I, Large (Klenow) Fragment (NEB, M0210), 0.8 U/µl of DNA, in 50 µl of 0.3 mM biotin-14-dATP (Thermo Fisher, 19524016), 0.3 mM dCTP/ dGTP/ dTTP for 1.5 hours at 37°C.

900 µl of ligation master mix were then add to the samples: 1.33x NEB T4 DNA ligase buffer (NEB, B0202), 1.1% Triton X-100, 1.33 mg of Bovine Serum Albumin and 2000 U of T4 DNA Ligase (NEB, M0202). Samples were incubated for 4 hours rotating. Nuclei were spun and resuspended in water and digested with proteinase K (Sigma, P6556) 1.2 mg/ml and 0.8% SDS at 55°C for 30 minutes. 130 µl of 5 M NaCl were added and samples were incubated at 65°C overnight. After ethanol precipitation, DNA was resuspended in 500 µl of 10 mM Tris-HCl pH8.0 and washed trice using the same buffer using Amicon filters (Millipore, UFC503096). Samples were diluted in 50 mM Tris pH 8.0, 0.1% SDS, 10 mM EDTA and sonicated to about 500 bp using a probe-based sonicator. Washes were then repeated as in the previous step. Biotinylated DNA was pulled down using 30 µl of 10 mg/ml Dynabeads MyOne

Streptavidin T1 beads (Life technologies, 65602) in 10 mM Tris-HCl (pH 7.5); 1 mM EDTA; 2 M NaCl for 15 minutes at room temperature rotating. Beads were washed in TWB: 5 mM Tris-HCl (pH 7.5); 0.5 mM EDTA; 1 M NaCl; 0.05% Tween 20 for 2 minutes at 55° C twice, resuspended in 20 µl 1× NEB T4 DNA ligase buffer, transferred to a new tube and incubated for 30 minutes at room temperature with 100 µl of: 1x NEB T4 DNA ligase buffer, 0.5 mM dNTPs, 50 U T4 PNK (NEB, M0201), 12 U T4 DNA polymerase I (NEB, M0203), 5 U DNA polymerase I, Large (Klenow) Fragment. After two washes in TWB at 55°C for 2 minutes, beads were transferred to 1× NEBuffer 2 (NEB, B7002S) and moved to a new tube. Beads were then incubated at 37°C for 30 minutes in 100 µl of 0.9X NEBuffer 2, 0.5 mM dATP and 25 U Klenow exo minus (NEB, M0212). Beads were washed again as before and transferred to T4 ligation buffer. DNA was ligated for 2.5 hours with Illumina adaptors in 50 µl of the following reaction mix: 1X T4 ligation buffer, 1.2 µM Illumina adaptors, 1U T4 DNA ligase. Beads were washed once again with TWB as before, to be then resuspended and stored in 50 µl of 10 mM Tris-HCl (pH8.0).

Libraries were amplified in six parallel 50 µl PCR reactions using Illumina primers and 4 µl of beads (or DNA removed off the beads by heating at 95°C for 20 min) per reaction after testing optimal amplification cycles. Size selection was performed on an agarose gel, isolating fragments between 300-700 base pairs, followed by purification with QIAquick Gel Extraction Kit (Qiagen 28704). Each library was sequenced on one lane of Illumina Nex-seq 500/550 (75 bp paired end reads). FASTQ files were processed with the distiller pipeline (https://github.com/mirnylab/distiller-nf, DOI:10.5281/zenodo.2630563) to obtain *.mcool* files used in downstream analyses. The original names of the cell lines used in these experiments, the name of the HiC raw files and the name used in the paper are: RFHF14 = het 14 = Rif1-FH; 14 tgWT A7 = tgWT_14A7 = Rif1-TgWT 1; 14 tgWT H4 = tgWT_14H4 = Rif1-TgWT 2; 14 tgwt G10 = tgWT_14G10 = Rif1-TgWT 3; 14 ΔP H1 = DP_14H1 = Rif1-ΔPP1 1; 14 ΔP H2 = DP_14H2 = Rif1-ΔPP1 2; 14 ΔP F8 = DP_14F8 = Rif1-ΔPP1 3; ESC B = WT_B = Rif1-WT 1; ESC F = WT_F = Rif1-WT 2; ESC 37.5 = WT_37.5 = Rif1-WT 3; ESC 24 = KO_24 = Rif1-KO 1; ESC 18 = KO_18 = Rif1-KO 2, ESC 28 = KO_28-2 = Rif1-KO 3.

**Protein extraction**

Cells were washed twice in cold DPBS and resuspended in hypotonic buffer: 25 mM Tris-HCl (pH 7.4); 50 mM KCl; 2 mM $MgCl_2$; 1 mM EDTA, proteinase inhibitor (Roche, 5056489001) at the concentration of $20\cdot10^6$ cells/ml. Samples were incubated for 20 minutes on ice, washed twice with hypotonic buffer and resuspended in the same volume of Benzonase buffer: 50 mM Tris-HCl (pH 8.0); 100 mM NaCl; 1.5 mM MgCl2; 10 % Glycerol; proteinase inhibitor (Roche, 5056489001). After 3 cycles of snap freezing and thawing, samples were supplemented with 50 U/ ml of benzonase (Sigma, E1014) and incubated at room temperature for 25 minutes. 0.2 % Triton X-100 was added and samples were incubated for 10 minutes at 4°C rotating. After centrifugation, supernatant was collected and quantified by Bradford (Bradford, 1976) (Biorad, 5000006). Immunoprecipitation was performed, SDS-Page and Western Blot analysis were performed as in (Foti et al., 2016).

**RNA extraction and qRT-PCR.**

Total RNA was extracted using RNeasy kit (QIAGEN, 74106) according to manufacturer's instructions. cDNA synthesis was performed using RevertAid H Minus reverse transcriptase (Thermo Scientific, K1632) and qPCR was performed using the SYBR Green reaction mix (Roche, 04887352001) on a LightCycler 96 Instrument. Gene expression data was normalized against GAPDH and the relative RNA expression levels were calculated using the Ct ($2^{\Delta\Delta Ct}$) method.

**3D-SIM and image analysis**

Cells were split on a coverslip on the 4$^{th}$ day of OHT treatment and after 5 hours after splitting were pulsed with 10μM BrdU for 15 minutes. After 2x washes in PBS at 37°C, cells were chased for 3 hours in $CO_2$ pre-equilibrated, medium at 37°C. After 2x washes in PBS, cells were fixed for 10 minutes in methanol free 3.7% formaldehyde. For immunostaining, fixed cells were permeabilised for 15 minutes in 0.5% TritonX-100/PBS, blocked for 30 minutes in 0.02% Tween 20/PBS/2% BSA, and stained with anti BrdU antibody (Biomol, Rockland, 600-401-C29, RRID:AB_10893609) for 60 min at 37 °C. Washes were performed in PBS/0.02% Tween 20. Immunostained cells mounted in Vectashield (Vector

Laboratories) were used for super resolution imaging. Samples were acquired on a 3D-SIM Deltavision OMX V3 microscope (General Electric) equipped with a 100 × 1.4 oil immersion objective UPlanSApo (Olympus), 405 nm, 488 nm and 593 nm diode lasers and Cascade II EMCCD cameras (Photometrics). After acquisition, the 3D-SIM raw data were first reconstructed and corrected for colour shifts with the help of the provided software softWoRx 6.0 Beta 19 (unreleased). Pseudo-widefield images were generated and exported during reconstruction too. In a second step, a custom-made macro in Fiji (Schindelin et al., 2012) finalised the channel alignment and established composite TIFF stacks, that were subsequently used for image analysis. For image analysis reconstructed super resolution wide field images were merged first with the corresponding pseudo-wide field stacks using FIJI. Pseudo-widefield channels were processed to enhance the focal pattern using the following steps: Difference of Gaussians filter with sigma1 = 10 and sigma2 = 1 pixels, followed by an automatic threshold (Otsu) and subsequent setting of all pixels below the threshold to 0. In contrast the super resolution replication foci were pre-segmented using the DAPI nuclear mask, followed by an automatic threshold (Triangle algorithm) and subsequently all pixels below the threshold were assigned to a value of 0, as described in detail in (Chagin et al., 2016). Further quantification was performed using Volocity 6.3 (Perkin Elmer). For the super resolution channel, replication foci were detected with the following settings: Find objects, histogram based segmentation with fixed lower threshold of 1 followed by a separate touching object steps with an object guide size of 0 µm3 followed by a filtering step to remove objects < 0.0002 µm3. The corresponding pseudo wide field foci were segmented using a fixed threshold of 1 and touching objects were separated with a guide size of 0.02 µm3. Finally, the pseudo wide filed foci were filtered to remove foci < 0.02 µm3. As a last step for each pseudo wide filed focus the overlapping super resolution foci (nano RFis) were calculated, based on the volume overlaps.

**Click-it reaction and Immunofluorescence**

mESCs were grown on gelatinized coverslips overnight. EdU (5-ethynyl-2´-deoxyuridine-Invitrogen A10044) was added to culture medium to 10 µM final concentration and incubated for 30 minutes.

After PBS washes, cells were pre-extracted with Triton buffer (0.5 % Triton X-100; 20 mM Hepes-KOH (pH7.9); 50 mM NaCl; 3 mM MgCl2; 300 mM Sucrose) for 2 minutes at 4°C before fixation with 3 % paraformaldehyde-2 % sucrose for 10 minutes at room temperature. Cells were permeablised with Triton buffer for 10 minutes at RT, followed by PBS washes. After short blocking with 3% BSA/PBS for 2 minutes, Click-it reaction cocktail (as in manufacturer's instructions- Flow Cytometry Assay Kit, C10424, Invitrogen) containing Alexa Fluor 647 (Invitrogen A10277) (or Flow Cytometry Assay Kit, C10425. Invitrogen) containing Alexa Fluor 488 azide (Invitrogen, A10266) was added to coverslips and incubated in dark for 30 minutes, followed by two washes with 3% BSA/PBS. Samples were blocked with PBG (0.2% (w/v) cold water fish gelatin-Sigma G-7765; 0.5% (w/v) BSA-Sigma A-2153, in PBS). Primary antibodies were diluted in PBG and then added to samples for 2-hours incubation at room temperature or 1 hour at 37C. After three PBG washes, secondary antibodies (Invitrogen) were diluted 1:800 in PBG and the incubation was in the dark for 45 min at room temperature. After PBS washes, coverslips were mounted with Vectashield with DAPI (Vector laboratories, H-1200). Images were acquired from Zeiss 880 Airyscan with 100x oil objective.

***Antibodies***

| Antigen | Source | Cat. # | Class | Use and dilution |
|---|---|---|---|---|
| HA | Biolegend | 901514 | Monoclonal Mouse | WB 1:1000<br>IF 1:3000<br>FACS 1:3000 |
| Rif1 | Buonomo et al. 2009 | 1240 | Polyclonal Rabbit | WB 1:3000 |
| SMC1 | Bethyl | A300-055A | Polyclonal Rabbit | WB 1:10000 |
| BrdU | Biomol (Rockland) | 600-401-C29 | Polyclonal Rabbit | IF 1:300 |
| MCM3 | Santa Cruz | sc-9850 | Polyclonal Goat | IF 1:200 |
| PP1$\alpha$ | Abcam | ab52619 | Polyclonal Rabbit | WB 1:1000 |

| | | | | |
|---|---|---|---|---|
| H3S10ph | Millipore | 06-570 | Polyclonal Rabbit | IF 1:400 |

***Primers***

| Primer | Sequence 5' – 3' | Reference |
|---|---|---|
| MERVL F | atgggtccaggaatcaaggg | This paper |
| MERVL R | gcctctggagccaaaacttc | This paper |
| GAPDH F | tgtgagggagatgctcagtg | This paper |
| GAPDH R | atggccttccgtgttcctac | This paper |
